# Multi-omics analysis of the mechanism of alfalfa and wheat-induced rumen flatulence in Xizang sheep

**DOI:** 10.1101/2024.12.04.626838

**Authors:** Tianzeng Song, Jing Wu, Xiaoming Zhang, Ayesha Khan, Jianzhao Cui, Khuram Shahzad, Gaofu Wang, Yangzong Zhaxi, Mingyan Shi, Wangsheng Zhao, Xunping Jiang, Long Li, Guiqiong Liu

## Abstract

This study induced a rumen flatulence model in Xizang sheep using alfalfa (HRF) and wheat grass (MRF) to explore related mechanisms. The pH of HRF and MRF groups was lower than the natural grass group. SCFA concentrations varied in different groups. Microbial analysis showed HRF and MRF groups had lower alpha and beta diversity and different compositions. Transcriptome analysis found different numbers of differentially expressed genes in each group. Spearman’s correlation analysis indicated Butyrivibrio was related to GLRX and DUOX2, and Quinella to several genes.

## 1 Introduction

Rumen flatulence is typically a fatal digestive system disease that primarily occurs in ruminants fed with leguminous plants or wheat pastures ^[1]^. Leguminous forages contain high concentrations of digestible proteins, which are released into the rumen and lead to the rapid proliferation of rumen microorganisms and fermentation, further resulting in the production of large amounts of gas in the rumen ^[2]^. The rapid proliferation of microorganisms, especially the formation of bacterial biofilms, leads to the production of excessive bacterial mucus ^[3]^, thereby increasing the viscosity of rumen contents and fermentation gases ^[4]^. Additionally, legumes also possess foaming properties due to glycosides (saponins), which can further promote flatulence ^[5]^. Howarth et al. demonstrated that high-protein forages mainly include alfalfa grass, clover grass, and wheat grass, which can promote the proliferation of certain harmful microbial populations in the rumen, thus affecting rumen health and inducing rumen bloating in ruminants ^[6]^.

Xizang sheep are one of the three original sheep breeds in China and are primarily distributed in the Qinghai-Tibet Plateau and surrounding areas ^[7]^. Having lived for generations at high altitudes and in low-oxygen environments, primarily through grazing, Xizang sheep are influenced by various factors that affect their rumen microorganisms. Long-term grazing can lead Xizang sheep to inadvertently consume certain grasses rich in protein and energy, predisposing them to rumen diseases.

Alfalfa is widely used among legumes due to its high nutritional value and yield. Studies have shown that sheep increase the viscosity of rumen contents after consuming fresh alfalfa grass ^[8]^. By comparing the rumen contents of sheep with and without bloat, it was found that the rumen contents of non-bloated sheep had higher mucus dissolution activity, while the rumen contents of bloated sheep had more foam with a darker color ^[9]^. Even after 24 hours of fasting, a small amount of foam remained in the rumen contents ^[10]^. Alfalfa is very important as livestock feed in both developed and developing countries because it contains a lot of protein and has a high dry matter yield ^[11]^. The main anti-nutritional component in this plant is saponins, which limit the nutrient absorption from alfalfa in ruminants ^[5]^. Lu et al. indicated that saponins in alfalfa may adversely affect the synthesis of rumen microbial proteins ^[12]^. Saponins cannot be effectively utilized by microorganisms, suggesting an antagonistic relationship between alfalfa saponins and microbial metabolism ^[5]^. Previous studies have shown that intragastric administration of alfalfa saponins can cause bloat in sheep, particularly at high concentrations, leading to a significant reduction in the number of rumen protozoa ^[13]^. Alfalfa saponins enter the stomach, promote the production of mucus by rumen bacteria, and alter the surface tension of the rumen. Therefore, they are considered as a contributing factor to the bloat formation ^[14]^.

Another study found that plant proteins are the main foaming agents responsible for bloating in ruminants. Plant proteins increase the viscosity of rumen contents, making it harder to expel gas as viscosity rises, which leads to foam formation. This foam can interfere with the belching process, resulting in bloat. Wheat forage contains a high proportion of plant protein and has low neutral detergent fiber contents ^[15]^. However, ruminants themselves lack the ability to degrade soluble proteins ^[16]^, so a large intake of high-concentration protein can impose a certain burden on the rumen.

Currently, there are few reports on rumen flatulence, and most studies remain focused on basic culture method or on targeted amplification and quantification ( qPCR ) of classical rumen bacteria ^[17]^. However, most of the microorganisms inhabiting the rumen ecosystem are unculturable or lack targeted genome amplification, and may therefore be overlooked as important contributors to the rumen foam and flatulence ^[18]^. Studies have shown that grazing increases six major bacterial groups associated with flatulence in the rumen contents of cattle in wheat pastures, indicating that foamy flatulence may be related to specific types of bacterial groups ^[19]^. Therefore, this study selected alfalfa and wheat forage to induce rumen flatulence model of Xizang sheep to simulate the ruminants with flatulence under natural conditions. To explore the mechanisms of action of host and microbiome and transcriptome.

## 2 Materials and Methods

### 2.1 Animal selection and experimental design

In this experiment, 24 healthy male Xizang sheep (provided by Longri Breeding Livestock Farm in Hongyuan County, Aba Prefecture, Sichuan Province) aged 7∼8 months, with a body weight of (21.2±25kg) were selected and divided into 3 groups using completely randomized experimental design. The control group (LRF) grazed freely on natural grass, while the treatment groups consisted of the high inflation group (HRF), which was fed alfalfa grass and the moderate inflation group (MRF), which was fed wheat grass. First, a 7-day pre-test was performed, followed by a 60 days formal test. The formal feeding time was August 15, 2023 (summer). All sheep were immunized using standardized procedures (deworming, vaccination, and ear marking). The treatment group was fed at 8:00 a.m. and 18:00 p.m. every day and was given free access to drinking water. On thefinal day of the experiment, Xizang sheep were fasted and withheld from water for 8 hours. Five Xizang sheep were randomly selected from the control group, and five Xizang sheep with the most obvious flatulence were selected from the treatment group for slaughter. Rumen contents and rumen epithelial tissues were collected. The samples were stored in DNase and RNase free freezer tube, immediately placed in liquid nitrogen, transported to the laboratory and stored at −80℃ for later use.

### 2.2 Experimental instruments

QuantStudio® 6 Flex fluorescence quantitative PCR instrument (Thermo Fisher Scientific, China), Qiagen Gel Extraction Kit (Qiagen, Germany), Applied Biosystems Agarose, Bio-Rad, Hercules, CA, USA; agarose gel imager (Beijing Wuzhou Technology Co., Ltd.), 9700 PCR instrument; agarose electrophoresis system Ready gas chromatograph (GC-2010 Plus; shimadzu, Kyoto, Japan); 721 spectrophotometer; centrifuge; gC-MS instrument, etc.

### 2.3 DNA extraction

The DNA from the swab samples was extracted using the PureLink™Microbiome DNA Purification Kit (Invitrogen, Carlsbad, CA) according to the manufacturer’s instructions. In brief, the swab samples were treated with Lysis Buffer and Lysis Enhancer, followed by vertexing to facilitate cracking. The supernatants were collected after centrifugation, and Cleanup Buffer was added and the mixture was immediately vortexed. A second Centrifugation was performed to remove any sediment. Binding Buffer was then added, the samples were loaded into spin column-tube assemblies, washed twice with Wash Buffer, and subsequently incubated with Elution Buffer. The resulting mixture was then centrifuged to collect the purified DNA into new sterile centrifuge tubes. DNA concentrations were measured using a NanoDrop spectrophotometer (Thermo Fisher Scientific, Inc., Waltham, MA, USA) and finally, the purified DNA samples were stored at −20℃.

### 2.4 16S rDNA amplicon sequencing and analysis

Sequencing of the 16S rDNA amplicon from fecal DNA of sheep was conducted by MetWare Biotechnology Co., Ltd. (Wuhan, China) using the Illumina NovaSeq 6000 platform (Illumina, San Diego, CA, USA) as previously reported ^[20]^. The raw data obtained were filtered and spliced to produce clean data. Denoising was performed using Deblur ^[21]^, generating Amplicon Sequence Variants (ASVs). The ASV sequences were annotated for species using Mothur (v1.48), and taxonomic information along with community compositions at various levels (phylum, class, order, family, genus, and species) were analyzed. The alpha diversity metrics, including Shannon, Simpson, Chao1, ACE, observed ASV, Goods coverage, and PD whole tree, were analyzed to assess species richness and evenness in the samples, as well as to identify common and unique ASVs across different samples. Beta diversity between groups was investigated using Principal Co-ordinates Analysis (PCoA), Principal Component Analysis (PCA), Non-Metric Multi-Dimensional Scaling (NMDS), and the Unweighted Pair-group Method with Arithmetic Means (UPGMA) based on Weighted UniFrac distance of ASV abundances. Differences in microbial composition and community structure of two groups were analyzed using the t-test and Wilcoxon test. Additionally, Tax4Fun2 (v1.1.5) was used for functional predictions of microbes. The Spearman correlation coefficient was applied to evaluate the relationship between microbes and DEGs, as well as inter-microbial relationships.

### 2.5 RNA extraction, cDNA synthesis and real-time quantitative PCR (qPCR)

Total RNA was extracted from rumen samples using RNAiso Plus reagent (Takara Biotechnology Co., Ltd., Dalian, China) according to the standard protocol. Subsequently, the mRNA was reverse transcribed into cDNA using the PrimeScript TM RT kit (Takara) containing gDNA Eraser. TB Green ® Premix Ex Taq TM II (Tli RNaseH Plus) was used for real-time quantitative PCR (qPCR) and performed in CFX96 real-time PCR detection system (Biorad, Hercules, CA, USA). The 2^-ΔΔCt^ technique was employed to determine the relative gene expressions.

### 2.6 RNA sequencing and analysis

RNA sequencing was performed by MetWare Biotechnology Co., Ltd. (Wuhan, China) using the Illumina NovaSeq 6000 platform (Illumina, San Diego, California, USA). The raw data was filtered to obtain clean data, which was then compared with the reference genome of Xizang sheep was replaced by Hisat2 ^[22]^ ftp://ftp.ensembl.org/pub/release-105/fasta/ovis_aries/dna/. Differentially expressed genes (DEGs) between the three groups were analyzed using FeatureCounts ^[23]^ and DESeq for comparison ^[24]^. The false discovery rate (FDR) was calculated using the Benjamini-Hochberg method to adjust the P-values. DEGs were identified based on |log2Fold Change|≥1 and FDR < 0.05. Data analysis, such as heat map generation, volcano plots, Gene Ontology (GO) and Kyoto Encyclopedia of Genes and Genomes (KEGG) pathways analyses, was performed using the Metware Cloud platform (https://cloud.metware.cn).

### 2.7 Statistical analyses

The data were expressed as mean ± standard error of mean (SEM) and analyzed using GraphPad Prism 8. An unpaired Student’s t-test was used to determine the differences between the two groups. A significance level of *p* < 0.05 was considered indicative of a significant difference, with * *p* < 0.05, ** *p* < 0.01, *** *p* < 0.001 in this study.

## 3 Results

### 3.1 Comparison of rumen fluid fermentation parameters among three groups of Xizang sheep

Changes in rumen PH values are shown in Fig. 1A, and rumen fermentation parameters are shown in Fig. 1B. The results showed that compared with the LRF group, the rumen PH values in the MRF and HRF groups were significantly reduced, with P-values 0.0063, 0.0003, respectively. Additionally, compared with LRF group, the concentrations of 4-MVA (*P* = 0.0186), 5-MCA (*P =* 0.0487), AA (*P* = 0.0493), BA (*P* = 0.0152), PA (*P* = 0.0219) and VA (*P* = 0.0207) were significantly increased in the MRF group. In the HRF group, the concentrations of 2-methylbutyric acid 2-BA (*P* = 0.0491), caproic acid CA (*P* = 0.0127), heptanoic acid HPA (*P* = 0.0008), isobutyric acid IBA (*P* = 0.0427), isovaleric acid IVA (*P* = 0.0497), octanoic acid OA (*P* = 0.0012), propionic acid PA (*P* = 0.0022) and valeric acid VA (*P* = 0.0004) were increased. Only the concentration of decanoic acid DEA (*P* = 0.0153) was significantly higher in the LRF group than in the other two groups, while the concentration of nonanoic acid NOA did not differ significantly among the three groups.

**Fig 1.**
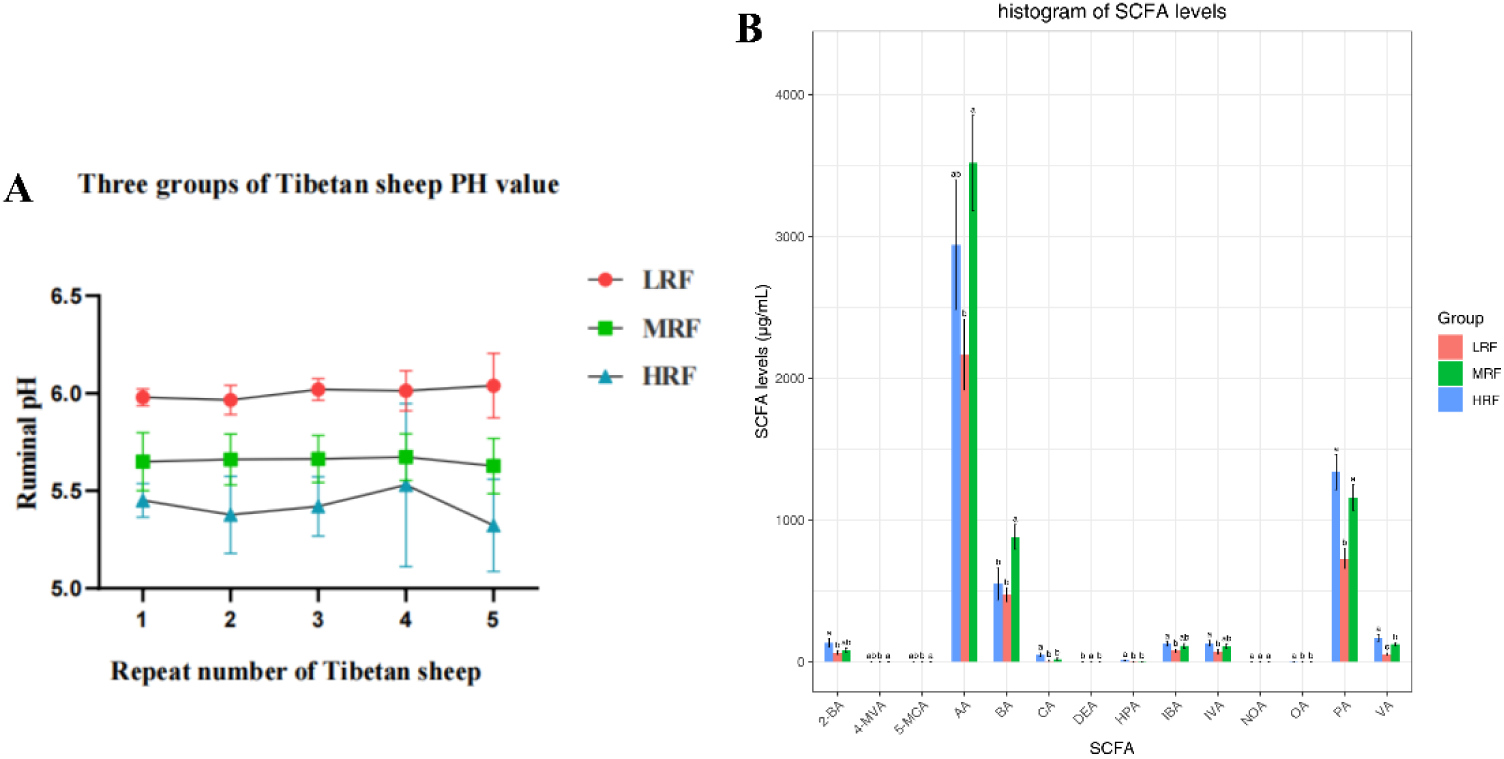
Comparison of rumen fermentation parameters of three groups of Xizang sheep. (A) Rumen PH. (B) short chain fatty acids.

### 3.2 Changes of α and β diversity of rumen microflora in Xizang sheep

The richness and diversity of the microbial community were mainly measured by Chao1 index, Shannon index, PD _ whole _ tree index and ACE index (Fig. 2ABCD). The α and β diversity of rumen microbiota varied with diet. The Chao1 index, Shannon index, PD _ whole _ tree and ACE index of rumen flora in Xizang sheep in LRF group were significantly higher than those in HRF group (*P*<0.01), but there was no significant difference between LRF group and MRF group. The Chao1 index, Shannon index, PD _ whole _ tree and ACE index of rumen microflora in MRF group were significantly higher than those in HRF group (*P*<0.05). It shows that the diversity of rumen flora in Xizang sheep is greatly affected by diet. As shown in fig. 2E, PCoA analysis showed that there were significant differences in β diversity of rumen microbiota among the three groups of Xizang sheep. The results showed that the abundance and diversity of rumen microflora of Xizang sheep in LRF group and MRF group were significantly higher than those in HRF group, indicating that the sensitivity of rumen flatulence induced by alfalfa was higher, and it was not conducive to the survival of microorganisms in the rumen environment with high flatulence.

**Fig 2.**
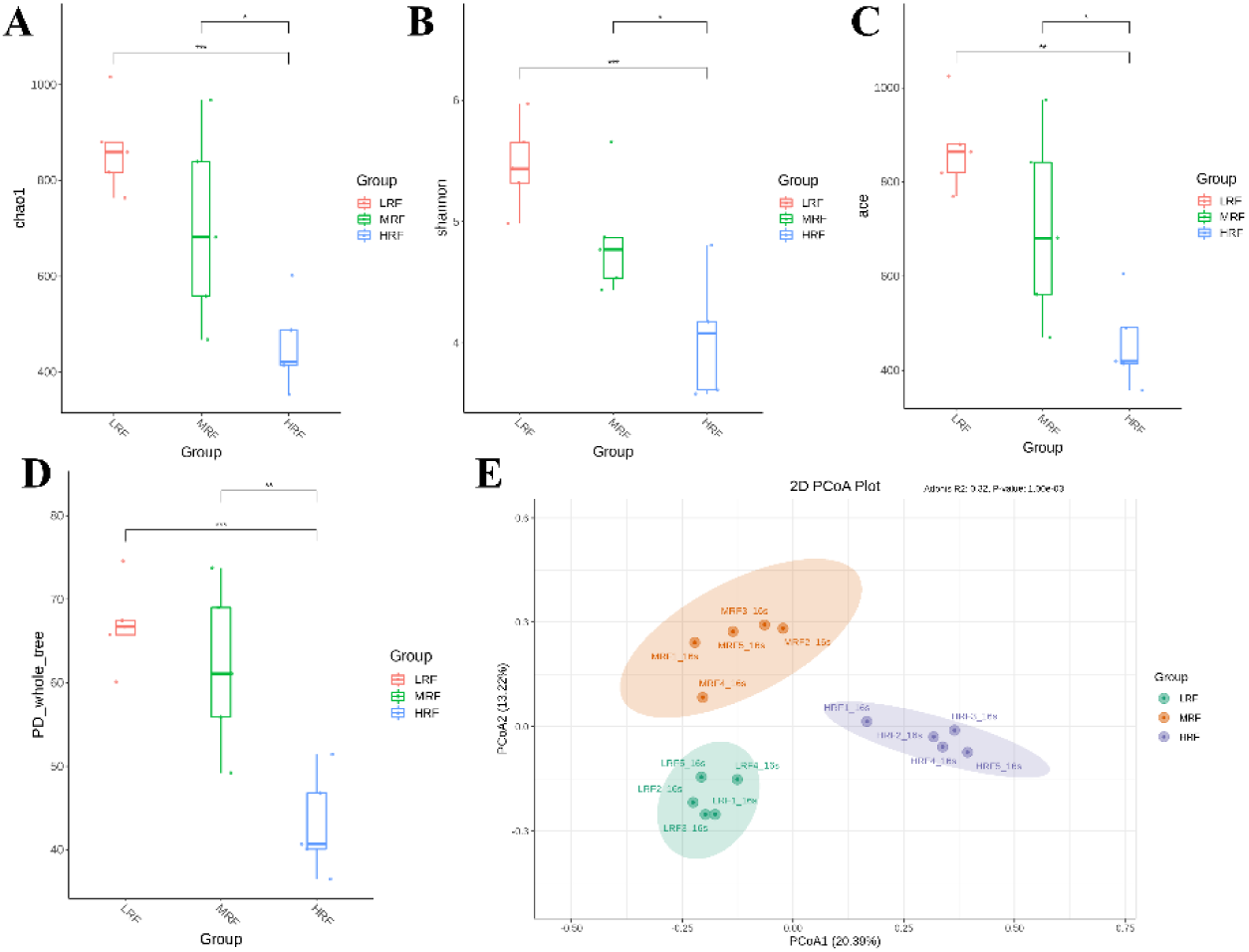
Comparison of α diversity changes and β diversity of rumen microbial communities among three groups of Tibetan sheep. (A) Chao1. (B) Shannon. (C) ACE. (D) PD-whole-tree. (E) PCoA. ⁎ denotes significant, ⁎⁎ denotes highly significant, ⁎⁎⁎ denotes extremely significant, LRF represents non-bloated Xizang sheep, MRF represents moderately bloated Xizang sheep, and HRF represents severely bloated Xizang sheep.

### 3.3 Changes of rumen microflora composition in Xizang sheep

At the microbial phylum level classification, the relative abundance of top 10 rumen microflora in Xizang sheep from the LRF, MRF and HRF groups was analyzed, as shown in Fig. 3A. Among these, *Bacteroidota* was he dominant phylum, accounting 61.0%, 47.5% and 72.4% in the three groups, respectively.. *Firmicutes* was the second most abundant, making up 35.5%, 36.6% and 24.2% in the three groups, respectively. Notably, the HRF group had the highest abundance of *Bacteroidota* and lowest abundance of *Firmicutes* among the groups, which are the two largest phyla of rumen microorganisms in Xizang sheep.

**Fig. 3.**
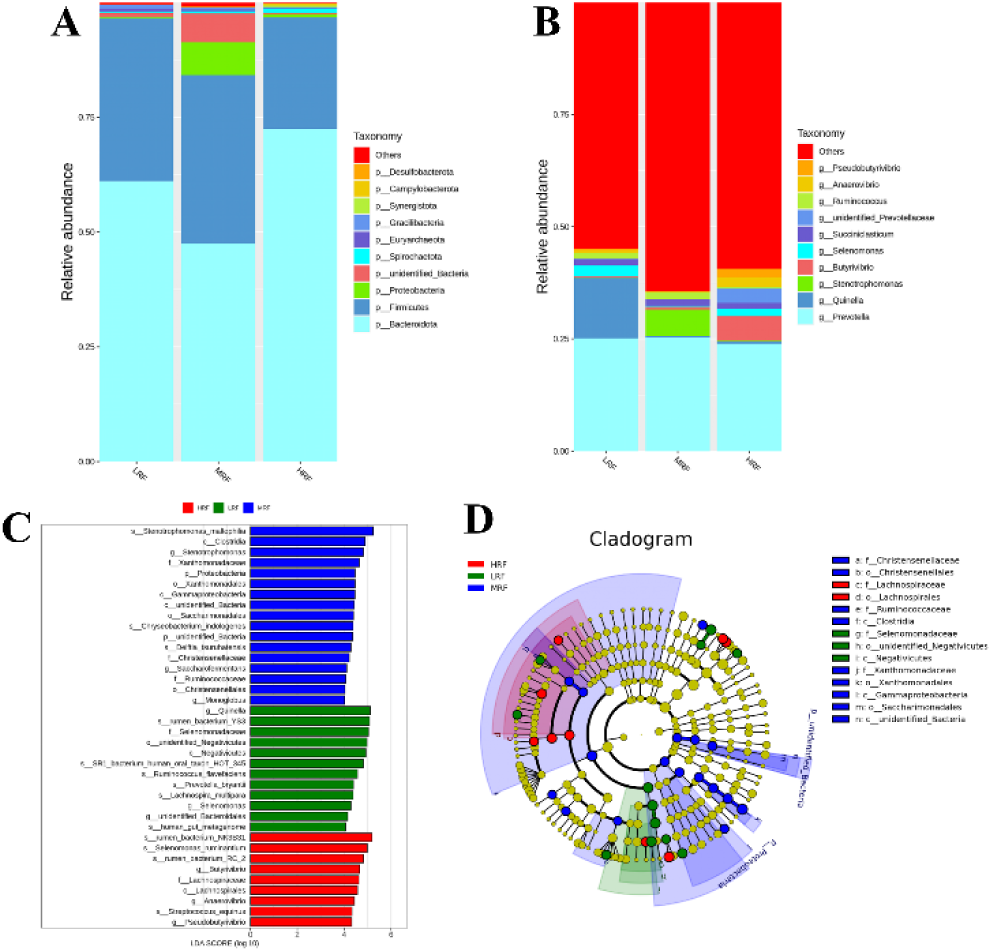
Variations in rumen microflora composition of Xizang sheep. (A) Composition of the microbial community at the phylum level and non-bloating groups. (B) Composition of the microbial community at the genus level and non-bloating groups. (C) Phylogenetic distribution from phylum to genus level of the microbial community, indicated by the branching diagram generated from LEfSe analysis. (D) LDA score histogram used to identify the differentially abundant bacterial genera between Xizang sheep and non-bloating groups.

At the microbial genus level, as shown in the Fig. 3B, *Prevotella* was common across all groups, with relative abundance of 25.1% in LRF, 25.3% in MRF and 23.9% in HRF. Additionally *Quinella* accounted for 13.4% of the dominant bacteria in the LRF group, but only 0.3% and 0.4% in the MRF and HRF groups, respectively. Stenotrophomonas was more dominant in the MRF group at 5.9% compared to 0.2% and 0.3% in the LRF and HRF groups, respectively. In contrast, *Butyrivibrio* was predominant in the HRF group at 5.4% with lower abundances of 0.2% and 0.6% in LRF and MRF, respectively.

LEfSe (LDA Effect Size) an analytical tool for biomarker discovery and interpretation across subgroups was employed to identify differentially enriched bacterial groups among the LRF, MRF, and HRF groups (Fig. 3C), using an LDA score threshold of 4 (Fig. 3D).This study identified 38 genera as key differentiators. Characteristic microflora in the MRF group included *Stenotrophomonas_maltophilia*, *Clostridia*, *Stenotrophomonas*, *Xanthomonadaceae*, *Proteobacteria*, *Xanthomonadales*, *Gammaproteobacteria*, *unidentified_Bacteria*, *Saccharimonadales*, *Chryseobacterium_indologenes*, *unidentified_Bacteria*, *Delftia_tsuruhatensis*, *Christensenellaceae*, *Saccharofermentans*, *Ruminococcaceae*, *Christensenellales* and *Monoglobus*. The LRF groupo was characterized by *Quinella*, *rumen_bacterium_YS3*, *Selenomonadaceae*, *unidentified_Negativicutes*, *Negativicutes*, *SR1_bacterium_human_oral_taxon_HOT_345*, *Ruminococcus_flavefaciens*, *Prevotella_bryantii*, *Lachnospira_multipara*, *Selenomonas*, *unidentified_Bacteroidales* and *human_gut_metagenome*. In the HRF group, the characteristics of microflora included *rumen_bacterium_NK3B31*, *Selenomonas_ruminantium*, *rumen_bacterium_RC_2*, *Butyrivibrio*, *Lachnospiraceae*, *Lachnospirales*, *Anaerovibrio*, *Streptococcus_equinus* and *Pseudobutyrivibrio*.

A hierarchical classification diagram from phylum to species level showed significant phylogenetic differences among the three groups. These findings suggest that dietary structure differences may lead to variations in rumen microflora composition among the groups, with significant enrichment of certain rumen microflora potentially contributing to rumen flatulence in Xizang sheep.

### 3.4 The construction and sequencing of the transcriptome library

Box plots and density plots display the variability and abundance of gene expression levels in individual samples (Fig. 4A, B) for the RNA sequencing analysis of 15 rumen epithelium specimens (LRF1, LRF2, LRF3, LRF4, LRF5, MRF1, MRF2, MRF3, MRF4, MRF5, HRF1, HRF2, HRF3, HRF4, HRF5).

**Fig 4.**
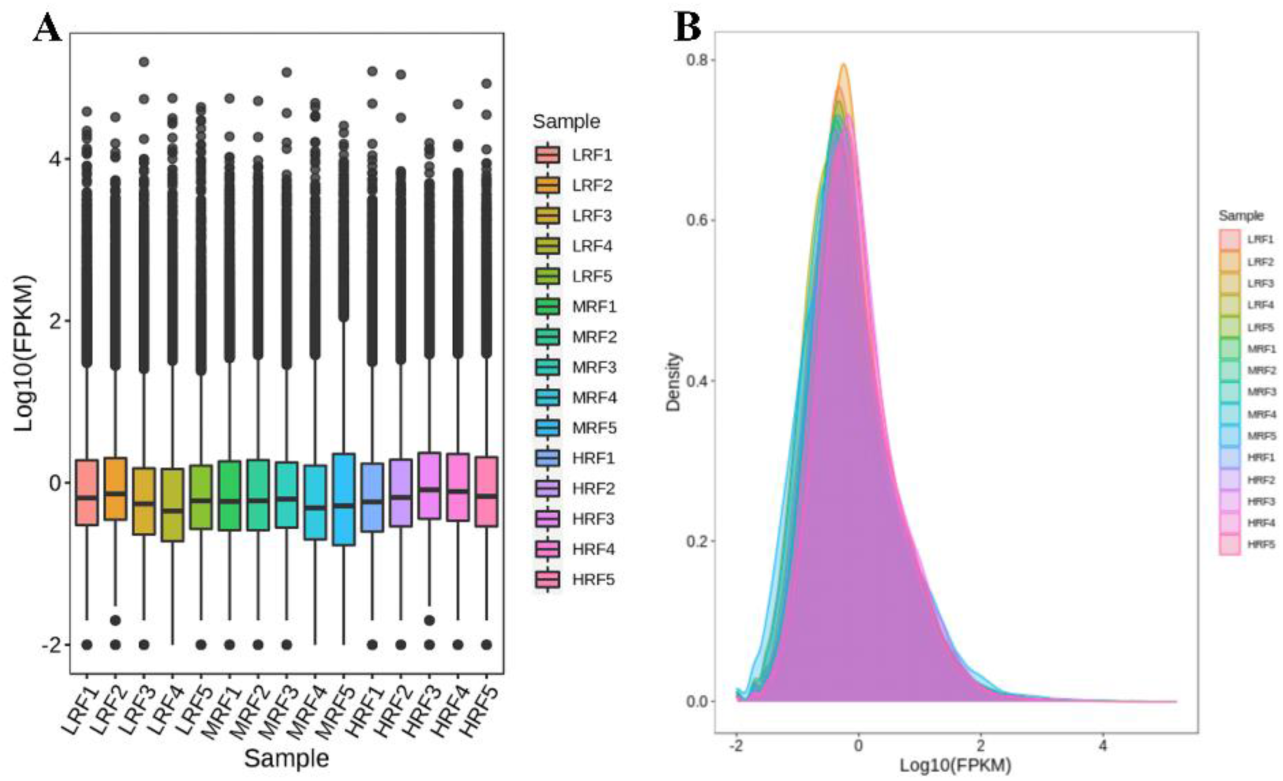
Differentially expressed genes (DEGs) among the three groups of Xizang sheep. (A) Overall gene expression levels and trends of expression in each group. (B) Clustering of DEGs among the groups.

### 3.5 Comparative Analysis of DEGs

#### Differentially Expressed Genes (DEGs) identified in the rumen epithelium of LRF and HRF

Further clustering analysis of the DEGs between the LRF and HRF groups revealed distinct clustering patterns of jhighly and lowly expressed genes in the rumen tissue samples. Heatmap analysis identified 859 shared DEGs between the LRF and HRF rumen epithelium samples (Fig. 5A, B). Specifically, compared to LRF, the HRF exhibited 348 upregulated and 511 downregulated DEGs. Gene Ontology (GO) enrichment analysis was performed to predict the functional properties of these DEGs (Fig. 5C). Results showed that DEGs observed in the rumen epithelium of LRF and HRF were enriched in biological processes (BP) including cellular process, metabolic process, response to stimulus, multicellular organismal process, and regulation of biological process, with significant differences between the two groups. In terms of cellular components (CC), the DEGs were involved in cellular anatomical entity and protein-containing complexes. For molecular functions (MF), DEGs were primarily concentrated in binding and catalytic activity. To explore the biological functions of the DEGs further, KEGG pathways analysis was conducted to assess pathway differences. As shown in Fig. 5D, the significantly enriched pathways included Carbon metabolism (2.23%) and Biosynthesis of amino acids (1.45%), with the Metabolism pathway showing the highest gene enrichment (9.95%).

**Fig 5.**
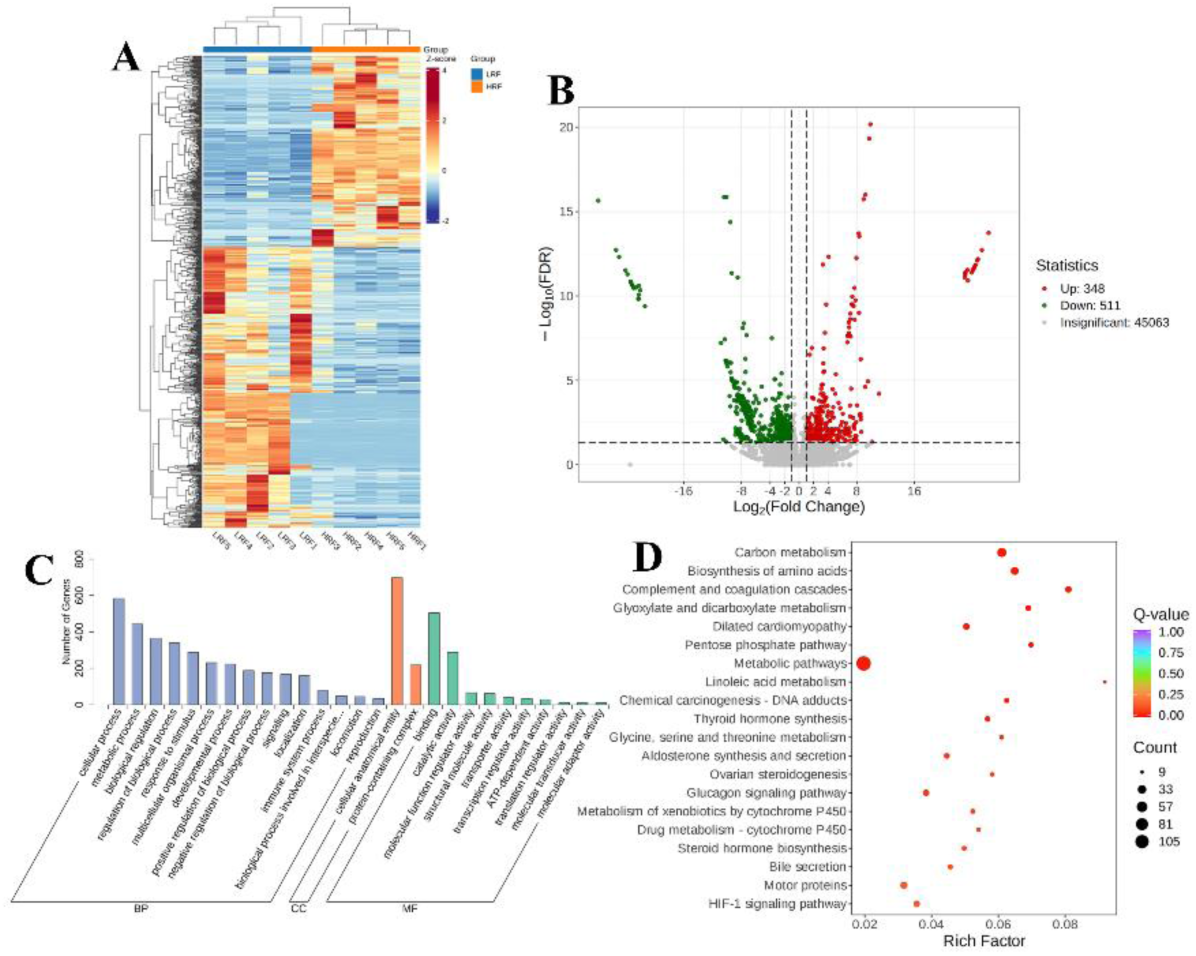
Differentially expressed genes (DEGs) in the rumen epithelium of Xizang sheep between LRF and HRF groups. (A) Clustering of DEGs. (B) Volcano plot of DEGs. (C) Gene ontology (GO) classification diagram. (D) KEGG enrichment plot of all the DEGs.

#### DEGs Identified in the rumen epithelium between LRF and MRF

Further clustering analysis of DEGs between the LRF and MRF groups (Fig. 6A), revealed significant clustering of highly and lowly expressed genes in the rumen tissue samples from each groups. Heat map diagram showed that there are 386 common DEGs between the LRF and MRF rumen epithelium samples (Fig. 6B). Specifically, MRF exhibited 201 upregulated and 185 downregulated DEGs compared to LRF group. GO functional enrichment analysis was conducted to identify the functions of DEGs in the rumen epithelium of Xizang sheep in the LRF and MRF groups, providing insight into their functional roles. As shown in Fig. 6C, the analysis was focused on biological processes (BP), cellular components (CC), and molecular functions (MF). In BP, DEGs were primarily involved in cellular processes, metabolic processes, responses to stimuli, multicellular organismal processes, and the regulation of biological processes. In CC, DEGs were involved in cellular anatomical entities and protein-containing complexes. In MF, DEGs were primarily concentrated in binding and catalytic activities. To determine the biological functions of DEGs, KEGG pathway enrichment analysis was performed to determine whether specific pathways were significantly enriched. As shown in Fig. 6D, the significantly enriched pathways included the cGMP-PKG signaling pathway (6.32%), fatty acid elongation (5.17%), biosynthesis of unsaturated fatty acids (4.6%), aldosterone synthesis and secretion (4.6%), ECM-receptor interaction (5.75%), adrenergic signaling in cardiomyocytes (5.17%), and the cAMP signaling pathway (5.17%). The pathways with the highest gene enrichment were metabolism and organismal systems.

**Fig 6.**
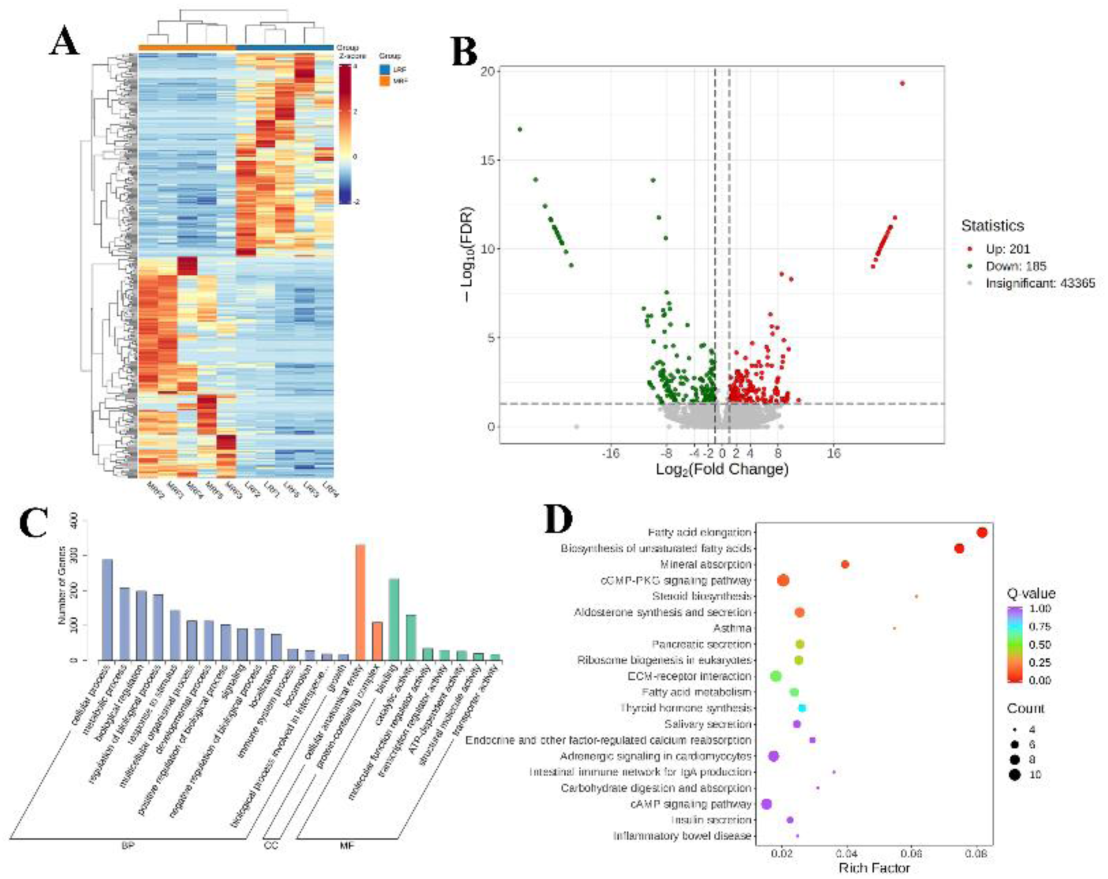
DEGs in the Rumen Epithelium of Xizang sheep between LRF and MRF Groups. (A) Clustering of DEGs. (B) Volcano plot. (C) Gene ontology (GO) classification diagram. (D) KEGG enrichment plot of all the DEGs.

#### DEGs identified in the rumen epithelium between HRF and MRF

Further clustering analysis of DEGs between the high-risk (HRF) and medium-risk (MRF) groups (Fig. 7A), revealed distinct clustering of high- and low-expressed genes in the rumen tissue samples from each group. Heatmap analysis identified 366 common DEGs between HRF and MRF rumen epithelium samples (Fig.7B). Specifically, the MRF group exhibited 128 upregulated and 238 downregulated DEGs compared to the HRF. To understand the functions of the DEGs, GO terms functional enrichment analysis was performed using the DEGs from the HRF and MRF groups in the rumen epithelium. As shown in Fig. 7C, the analysis focused on BP, CC, and MF. Within BP category, DEGs were mainly involved in cellular processes, metabolic processes, responses to stimuli, multicellular organismal processes, regulation of biological processes, and biological regulation. In the CC category, DEGs were involved in cellular anatomical entities and protein-containing complexes. In the MF category, DEGs were primarily concentrated in binding and catalytic activities. To determine the biological functions of the DEGs, KEGG pathways enrichment analysis was conducted to identify any significantly enriched pathways. As indicated by Fig. 7D, significantly enriched pathways included dilated cardiomyopathy (5.52%), ECM-receptor interaction (6.63%), motor proteins (6.63%), protein processing in the endoplasmic reticulum (7.18%), microRNAs in cancer (6.63%), adrenergic signaling in cardiomyocytes (5.17%), focal adhesion (8.29%), PI3K-Akt signaling pathway (9.39%), MAPK signaling pathway (7.73%), and proteoglycans in cancer (6.63%). The pathway with the highest gene enrichment was Human Diseases.

**Fig 7.**
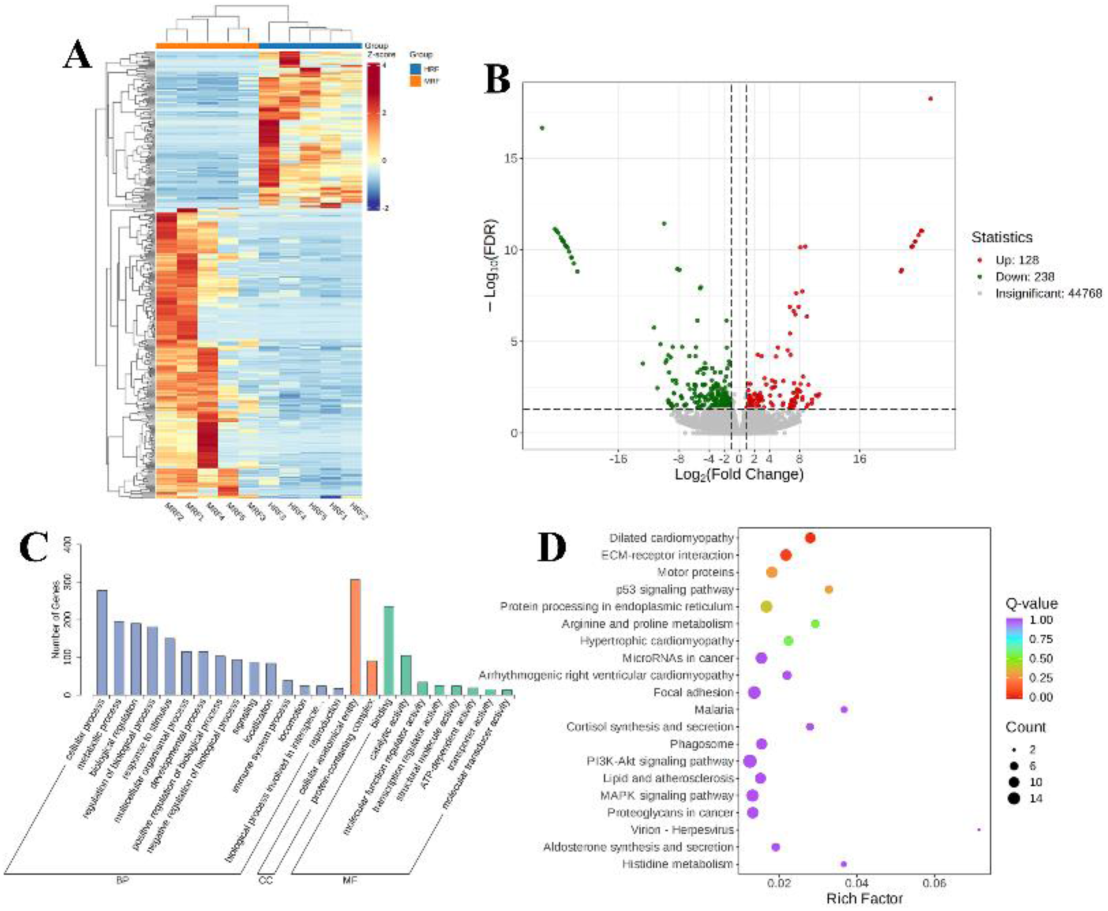
DEGs in the Rumen Epithelium of Xizang sheep between LRF and MRF Groups. (A) Clustering of DEGs. (B) Volcano plot. (C) Gene ontology (GO) classification diagram. (D) KEGG enrichment plot of all the DEGs.

### 3.6 Correlation analysis between rumen microbiota and differentially expressed genes

The Spearman correlation heatmap was used to analyze the relationship between rumen microbes and DEGs in the rumen to explore the effects of different diets on the host. As shown in Fig. 8, we observed a significant negative correlation between the *Butyrivibrio* microbiota and the *GLRX* and *DUOX2* genes. *Quinella* was significantly negatively correlated with *PI3*, *GLRX*, *SFTPC* and *CLDN7* genes, but positively correlated with *CP* and *IGFBPI*. Therefore, it can be inferred that the most affected species included *Butyrivibrio* and *Quinella*, which were found as the dominant genera in the HRF and LRF groups, respectively.

**Fig 8.**
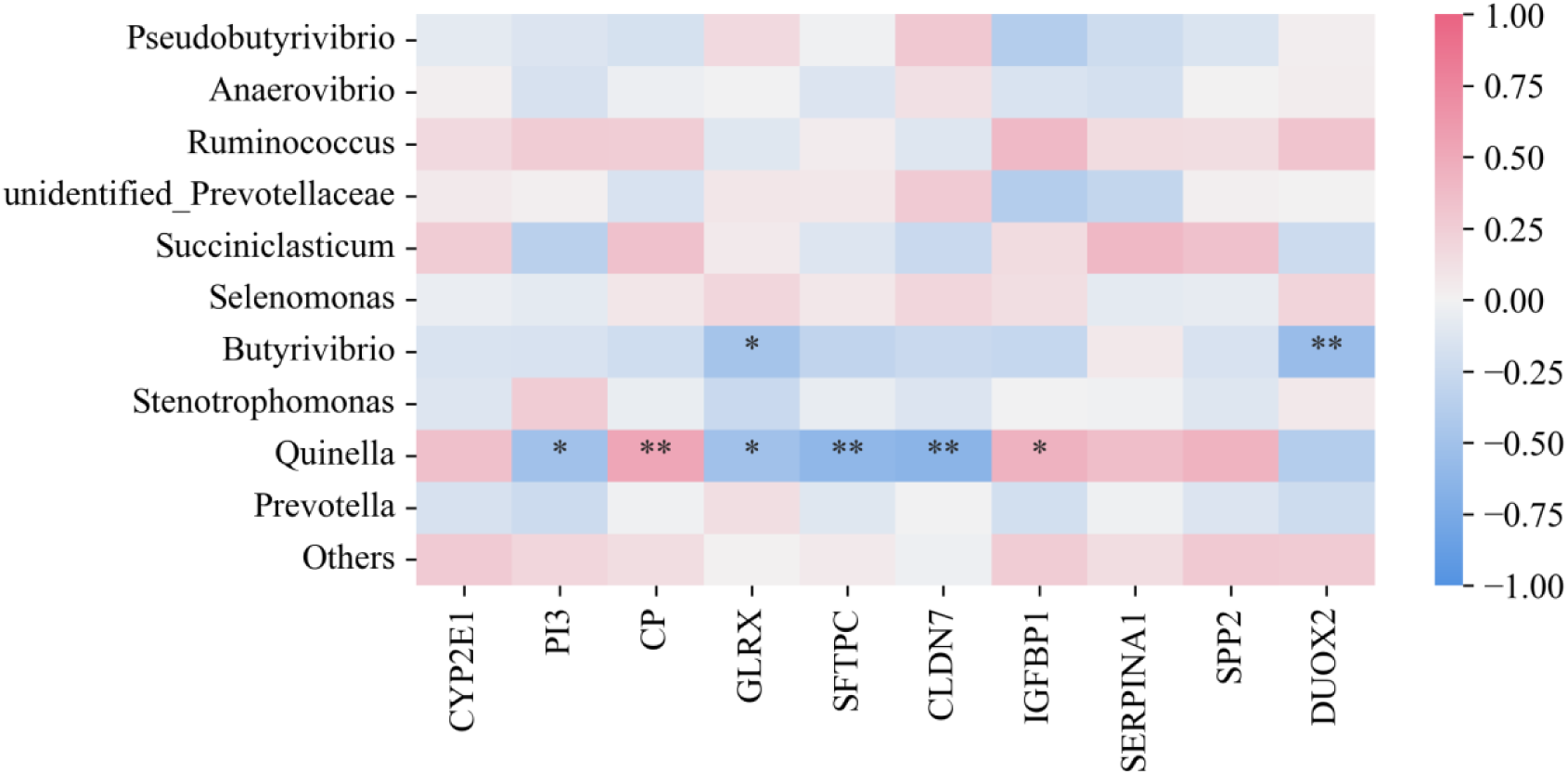
Correlation analysis between rumen microbiota and differentially expressed genes. ⁎denotes significant, ⁎⁎denotes highly significant.

### 3.7 RT-qPCR validation analysis

To confirm the reproducibility and reliability of gene expression levels in the RNA sequencing analysis, six randomly selected DEGs (*CYP2E1*, *PI3*, *IGFBP1*, *SPP2*, *DUOX2*, and *CP*) were validated using qRT-PCR. The results were consistent with the RNA sequencing findings (Fig. 9).

**Fig 9.**
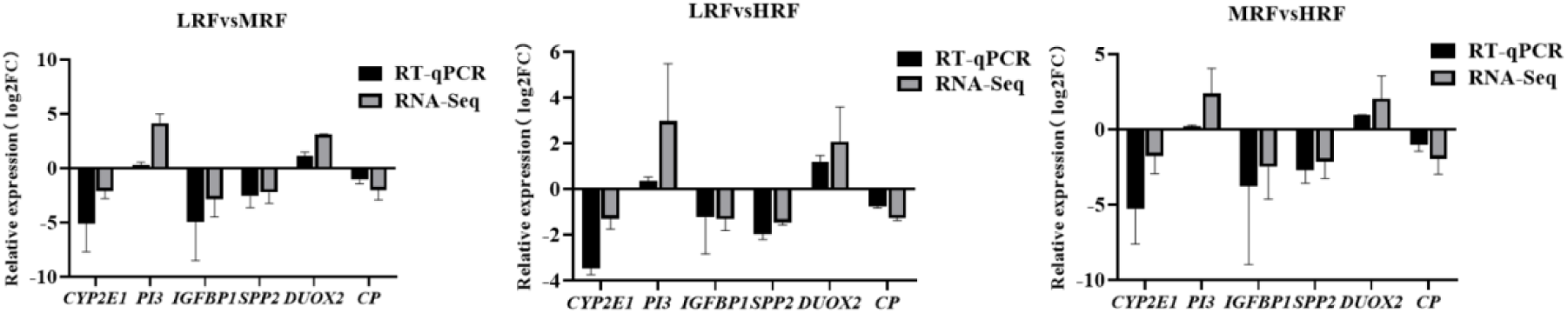
Verification of differentially expressed genes.

## 4. Discussion

In recent years, the development of various omics technologies has enabled researchers to explore more deeply of the changes between hosts and microorganisms. Numerous studies have demonstrated that the composition and function of gastrointestinal microbiota can influence health and disease, as well as affect metabolism, immunity, and neurobehavior ^[25]^. Other studies have indicated that certain genes are associated with disease and immunity, highlighting their significant roles in maintaining health ^[26]^. Therefore, this study extensively employs analytical techniques such as rumen transcriptomics and microbiology to investigate the interactions between the host and microbiota in the rumen of Xizang sheep, as well as the impact of rumen bloat on the host and microbiota. The aim is to further explore the regulatory mechanisms underlying bloat disorders.

The study identifies wheat grass, alfalfa, and oat grass as major annual forages on the Tibetan Plateau, characterized by drought resistance, cold tolerance, strong adaptability, high forage yield, and high nutritional value, making them well-suited for the harsh high-altitude environment ^[27, 28]^. These forages have become important feed for livestock, particularly as supplementary feed during the winter ^[29, 30]^. Research indicates that when the level of wheat hay decreases, the protein content increases, while the digestibility of the feed significantly declines ^[31]^. At this point, ruminants may not fully absorb and utilize the protein, and undigested protein may undergo abnormal fermentation in the gastrointestinal tract, producing harmful metabolic by-products that can damage gastrointestinal health and increase the risk of gastrointestinal diseases. In addition, studies have shown that feeding barley can lead to liver damage in cattle ^[32]^. Previous research has also indicated that high protein content is associated with diseases such as bloat and indigestion in ruminants. Their experiments demonstrated that animals fed a low-fiber diet experienced bloating or digestive disorders during the trial period ^[33]^. Our study corroborates these findings. In this research, we successfully induced rumen bloat in Xizang sheep using wheat grass and alfalfa, and analyzed the causes of rumen bloat by comparing the diversity of rumen microbiota and the differentially expressed genes in the rumen epithelium among different groups.

The rumen is an essential digestive organ in ruminants, and its internal environment can be easily influenced by the composition of the diet. Changes in dietary structure can lead to alterations in ruminal pH and volatile fatty acids (VFAs) ^[34]^. In this study, the pH measurements of three groups of Xizang sheep revealed that the pH of the rumen in bloat-induced sheep fed wheat and alfalfa was significantly lower than that of non-bloat sheep, which is consistent with the results of ruminal acidosis ^[35]^, Previous studies have also shown that the rumen pH significantly decreases in goats with bloat induced by high-concentrate diets ^[36]^. Therefore, it can be inferred that rumen diseases may lead to changes in pH levels.The accumulation of VFAs in the rumen can increase osmotic pressure, potentially damaging the ruminal epithelial cells and increasing the risk of pathogenic substances transferring into the bloodstream, thereby affecting metabolism and overall health ^[37]^. The dietary components that disrupt the rumen environment are largely determined by the levels of protein and fiber ^[38]^. The diet can influence the fermentation patterns of the rumen in ruminants. Studies have shown that volatile fatty acids, particularly acetate, propionate, and butyrate, are the primary metabolic products generated during the anaerobic fermentation process carried out by ruminal microbes, providing 70% to 80% of the energy required for the vital activities of ruminants ^[39]^. Additionally, the concentration of VFAs is an important indicator of ruminal fermentation efficiency. There are significant differences in dietary composition between grazing and housed systems ^[40]^, pasture-based diets typically have higher crude protein (CP) and soluble carbohydrate levels, with lower starch content ^[41]^. In our experiment, the concentrations of acetate, propionate, and butyrate in the ruminal contents of bloat-induced Xizang sheep fed wheat and alfalfa were significantly higher than those in non-bloat sheep, indicating that different feeds have a notable impact on ruminal fermentation in Xizang sheep. This is likely due to the fiber components and structural carbohydrates in wheat, which typically increase the levels of acetate, propionate, and butyrate ^[42]^. Another study noted that as the proportion of concentrate in the diet increases, the level of butyrate in the rumen also rises ^[43]^. Alfalfa contains a large amount of protein, and the high protein content can influence the fermentation patterns in the rumen. Furthermore, the non-structural carbohydrates in alfalfa are easily fermented by rumen microbes, leading to the production of increased amounts of acetic acid, butyric acid, and propionic acid ^[44]^.

The rumen hosts a large population of microorganisms, including bacteria, fungi, and protozoa, which together form a complex ecosystem ^[45]^. Alpha diversity can be used to assess the diversity of bacteria in the rumen of ruminants, with the Shannon index reflecting the evenness and diversity of rumen bacteria ^[46]^. Generally, a higher Shannon index indicates greater bacterial diversity within the sample. A larger PD_Whole_tree index and Chao1 index signify higher bacterial abundance in the sample ^[47]^. In this study, the Shannon index, PD_Whole_tree index, and Chao1 index showed highly significant differences among the three groups, with the MRF group and HRF group significantly lower than the LRF group. Additionally, PCoA analysis revealed significant differences in the beta diversity of the rumen microbial communities among the three groups of Xizang sheep. Previous studies found that the Shannon index, PD_Whole_tree index, and Chao1 index of rumen bacteria were significantly lower in goats with rumen bloat induced by high-concentrate diets compared to non-bloat goats, consistent with the results of this study, indicating that rumen bloat has a significant impact on the diversity of the rumen microbial community in Xizang sheep.

The composition and structure of rumen bacteria are critical for maintaining homeostasis, enhancing nutrient digestion and absorption, and supporting overall health ^[48]^. The ratio of Firmicutes to Bacteroidetes has been used as a key indicator to evaluate how microorganisms affect the host’s energy needs ^[49]^. Firmicutes play a major role in the degradation of fibrous materials, while Bacteroidetes are primarily responsible for the breakdown of non-fibrous substances ^[50]^. Both Bacteroidetes and Firmicutes are major microbial groups in the rumen and have a mutualistic symbiotic relationship that assists the host in energy absorption or storage. Therefore, the balance between Bacteroidetes and Firmicutes in the gastrointestinal tract is crucial for the fermentation of polysaccharides, and their ratio is significant; a reduction in Bacteroidetes or an increase in Firmicutes can promote obesity ^[51]^. Moreover, high-fat diets have been shown to reduce mucin in the intestines of rats, decrease microbial diversity, and weaken the intestinal barrier, increasing the risk of pathogenic bacteria such as Firmicutes invading and decreasing gut barrier-repairing bacteria like Bacteroidetes ^[52]^. This study shows that Bacteroidetes and Firmicutes are the dominant phyla in the three groups of Xizang sheep, providing the host with more energy to adapt to the high-altitude habitat. However, the abundance of Bacteroidetes in the MRF group is significantly lower than in the other two groups (*P*<0.01), while the abundance of Firmicutes is significantly higher than in the LRF and HRF groups (*P*<0.01). This may be due to limited capacity for nutrient absorption from wheat in the rumen of Xizang sheep. The occurrence of ruminal bloat varies among individual ruminants, depending on the fermentation rate of wheat forage, the production and passage of ruminal gases, and the foaming characteristics of ruminal contents ^[53, 54]^. The phylum Proteobacteria and unidentified Bacteria are dominant in the MRF group, which may be attributed to specific substances in wheat forage that promote the growth of these bacterial phyla. Additionally, the abundance of unidentified Bacteria is particularly high in the MRF group (*P*<0.01), suggesting that the fermentation characteristics of wheat feed lead to significant differences in the rumen microbial community.

*Prevotella* is generally regarded as a dominant genus in the rumen microbiome ^[55]^. In this study, the genus *Prevotella* was identified as the dominant genus across all three groups, while *Quinella* was the dominant genus in the LRF group, Stenotrophomonas in the MRF group, and *Butyrivibrio* in the HRF group. *Prevotella* plays a central role in carbohydrate and hydrogen metabolism, and a high abundance of *Prevotella* in ruminants is associated with a healthy microbiome ^[56]^. The capacity of *Prevotella* to ferment carbohydrates is also reflected in the productivity of cattle. It has been reported that, in a study involving a group of lactating Holstein cows, animals with higher protein and fat content in their milk exhibited higher concentrations of acetate, butyrate, and propionate in their rumen fluid ^[57]^. Furthermore, methane production was positively correlated with VFA concentrations (primarily acetate and butyrate), while rumen pH significantly decreased, which aligns with the findings of this study.

In this study, the genus *Prevotella* was identified as the dominant genus across all groups, with *Quinella* emerging as the dominant genus in the LRF group. The abundance of methanogenic archaea is proportional to the occurrence of rumen bloat. *Quinella*, a hallmark rumen bacterium first reported in 1913, has been shown to increase in number in the rumen of sheep, correlating with lower methane (CH₄) emissions ^[58]^. The fact that *Quinella* is the dominant genus in the LRF group, which is less sensitive to bloat, suggests that *Quinella* may play a role in alleviating the occurrence of bloat. However, how this genus contributes to the lower methane emissions in these sheep remains to be investigated. Previous research has indicated that *Stenotrophomonas* is associated with inflammation and tends to increase in the rumen of cows suffering from subacute ruminal acidosis ^[59]^. Furthermore, administering *Stenotrophomonas* to lactating mice can induce mastitis. Wheat-based diets have also been linked to health issues in cows due to their high dry matter content, leading to increased levels of *Stenotrophomonas* in their rumen, which positively correlates with acetate, propionate, and total volatile fatty acid (TVFA) levels. Thus, it can be inferred that the high abundance of *Stenotrophomonas* in the MRF group of Xizang sheep subjected to wheat feeding was a result of inflammation induced by the diet, contributing to rumen disorders. Members of the genus *Butyrivibrio* are significant components of the rumen microbiota ^[60]^. Early characterization of *Butyrivibrio* has shown that some strains possess the ability to degrade cellulose, while most can metabolize xylan and pectin substrates. It has been reported that ruminal *Butyrivibrio* can utilize various soluble and some insoluble substrates, fermenting carbohydrates into butyrate, formate, lactate, and acetate ^[61]^. Additionally, *Butyrivibrio* exhibits metabolic versatility and can utilize various insoluble substrates, although it is not classified as a cellulose-degrading bacterium. Studies have indicated that while *Butyrivibrio* hungatei can use oligosaccharides and monosaccharides as growth substrates, it cannot hydrolyze proteins or degrade fibers ^[62]^. Therefore, it is hypothesized that the high protein content in alfalfa stimulates substantial *Butyrivibrio* production in the rumen of the HRF group, leading to extensive fermentation of protein and gas production, thereby promoting the occurrence of bloat in Xizang sheep. Overall, the analysis of rumen microbiota in this study indicates that different dietary regimens affect the microbial composition in the rumen of Xizang sheep. These specific microbes can contribute to the development of rumen disorders such as bloat. The identification of dominant genera within each group illustrates their potential role in the occurrence of rumen bloat in Xizang sheep.

Rumen bloat involves a series of encoded genes, and to investigate the regulatory gene changes among the LRF, MRF, and HRF groups, an RNA-seq analysis was conducted. The results revealed a total of 1417 differentially expressed genes (DEGs) across the three groups. Specifically, between the LRF and HRF groups, there were 859 shared DEGs, with 511 downregulated and 348 upregulated; between the LRF and MRF groups, there were 386 shared DEGs, with 185 downregulated and 201 upregulated; and between the HRF and MRF groups, there were 366 DEGs, with 238 downregulated and 128 upregulated. The functional annotation of these DEGs indicated their involvement in cellular processes, metabolic processes, responses to stimuli, multicellular organism processes, biological regulation, cellular component organization, protein complexes, binding, and catalytic activity regulation. By integrating the rumen microbiome with host gene expression patterns, we found correlations between certain rumen microbes and host genes. Research suggests that gene expression may play a mediating role between microbial communities and host functionality. For instance, *CYP2E1* is involved in disease occurrence and immune-mediated mechanisms, with downregulation of *CYP2E1* observed in nearly all immune-mediated diseases, making it a significant factor in disease development ^[63]^. Consequently, downregulation of *CYP2E1* may impair animal health. *CYP2E1* is a critical enzyme in the metabolic activation of various toxic substances, including nitrosamines, benzene, vinyl chloride, and halogenated solvents like trichloroethylene ^[64]^. Cows exhibiting signs of ketosis show abnormal expression of *CYP2E1* compared to healthy cows. The oxidation of multiple substrates mediated by *CYP2E1* can release substantial reactive oxygen species, leading to lipid peroxidation and cytotoxicity ^[65]^. Therefore, *CYP2E1* serves as a significant marker involved in disease development. The *PI3* signaling pathway is implicated in gastrointestinal regulation. Studies found that gas in the gastrointestinal tract may result from the activation of the PI3K/AKT signaling pathway, with the production of phosphatidylinositol 3-kinase being triggered during such conditions ^[66]^. Abnormal activation of the PI3K signaling pathway has been observed across various diseases ^[67]^. In this study, the upregulation of *PI3* expression was related to the PI3K/AKT pathway, suggesting that bloat in ruminants may lead to elevated *PI3* expression. When there is an excess of growth hormone or metabolic disorders, levels of *IGFBP1* decrease, potentially resulting in severe physiological phenomena such as abdominal spasms, gastrointestinal bloat, and diarrhea ^[68]^. In this study, *IGFBP1* showed a downregulated trend, indicating a potential relationship between gastrointestinal bloat and altered *IGFBP1* levels. Previous research found that the expression of *IGFBP5* did not correlate with the growth and proliferation of rumen papillae in Holstein calves, but *IGFBP5* in rumen papillae indicated subacute acidosis, damaging the gastric epithelial cells in Holsteins ^[69]^. Differential expression of *IGFBP3* and *IGFBP6* in non-lactating Holstein cows was reduced due to grain-induced subacute acidosis, resulting in increased concentrations of myocardial lactate and LPS, which harmed gastric epithelial cells ^[70]^. Secreted phosphoprotein 2 (*SPP2*) is primarily produced in the liver and transported to other tissues, where it has been identified as a significant bone matrix protein involved in the regulation of bone metabolism ^[71]^. *SPP2* may interact with insulin-like growth factor binding protein 1 (*IGFBP1*), potentially inhibiting the progression of liver fibrosis and cirrhosis ^[72]^. Previous studies have also reported a consistent and significant decrease in *SPP2* expression in tumors, with reduced *SPP2* levels in hepatocellular carcinoma (HCC) being associated with advanced clinical and pathological features, indicating a potential tumor-suppressive role for *SPP2* in HCC ^[73]^. Research has demonstrated that *SPP2* can inhibit tumor growth and induce tumor apoptosis by eliminating the pro-tumor functions of bone morphogenetic protein 2 (*BMP-2*) signaling ^[74]^. These findings suggest that *SPP2* and related genes may play a role in disease development, leading to adverse effects in Xizang sheep that are particularly sensitive to bloat.

Through the analysis of the rumen microbiome and host gene expression patterns, we identified multiple associations between differentially expressed rumen epithelial genes in LRF, MRF, and HRF groups of Xizang sheep and the rumen bacteria. Evidence for the relationship between host gene expression and the composition of the rumen microbiome suggests that gene expression may mediate interactions between microbial communities and host functions ^[75]^. In this study, a significant negative correlation was found between the genus *Butyrivibrio* and the *GLRX* and *DUOX2* genes. In addition, *Quinella* was negatively correlated with genes *PI3*, *GLRX*, *SFTPC* and *CLDN7*. It was significantly positively correlated with *CP* and *IGFBP1* genes. Previous research has indicated that reduced *GLRX* expression exacerbates liver fibrosis, with observed decreases in *GLRX* and increases in PSSG in fibrotic mice and human liver samples ^[76]^. Furthermore, *GLRX1* deficiency has been shown to make male mice sensitive to high-fat diets ^[77]^, with *GLRX1* deficiency worsening liver damage and oxidative stress induced by high-fat diets ^[78]^. An increase in *DUOX2* expression affects intestinal immune homeostasis in mice, and mutations in *DUOX2* can lead to hypothyroidism ^[79]^. Notably, Butyrivibrio is an important degrader and user of lignocellulosic plant materials ^[80]^. These results suggest that the sensitivity to bloat in Xizang sheep may be influenced by the interplay of various microbes and differentially expressed host genes.

## 5. Conclusions

In summary, this study revealed the changes of volatile fatty acids, rumen microbiota and rumen transcriptome in Xizang sheep with strong flatulence sensitivity. We found that PH, PA, and VA were significantly higher in the MRF and HRF groups than in the LRF group. In addition, changes in Butyrivibribrio and Quinella may be the cause of rumen flatulence in Xizang sheep. Genes related to immunity and metabolism were also found to be significantly down-regulated. In conclusion, our study provides insight into the potential interactions between host genes and rumen microbiota in the rumen of Xizang sheep.

## Data Availability Statement

All data used in this study are available upon request from the corresponding or first author. In addition, the dataset created specifically for this study has been deposited in the NCBI Sequence Read Archive, which can be found under the BioProject identifier PRJNA1178088 and PRJNA1180951.

## Author Contributions

T.S. and J.W. were in charge of collecting and analysing the samples as well as drafting and editing the initial draft. X.Z. and A.K. was in charge of experiment design and conceptualisation. The project data processing was handled by J.C. and K.S. The editing of the paper was assigned to G.W. and Y.Z. The integration and review were handled by M.S., X.J. and L.L. The management of the project, as well as the utilisation and management of finances, fell to G.L. and W.Z. After reading the published version of the manuscript, all writers have given their approval.

## Institutional Review Board Statement

This study adheres to animal ethics principles, and all experiments have been approved by the Livestock and Poultry Breeding Committee of Southwest University of Science and Technology (SWUST), with the sample collection protocol authorized (Approval No. L2022014). To maximize the welfare and rights of the animals, all animal feeding experiments and related sample collections in this study were conducted in strict accordance with relevant regulations and requirements.

## Informed Consent Statement

Informed consent was obtained from animal owners for all animal husbandry procedures described in this study.

